# Inwardly rectifying potassium channels regulate membrane potential polarization and direction sensing during neutrophil chemotaxis

**DOI:** 10.1101/2025.03.06.641746

**Authors:** Tianqi Wang, Daniel H Kim, Chang Ding, Dingxun Wang, Weiwei Zhang, Martin Silic, Xi Cheng, Kunming Shao, TingHsuan Ku, Conwy Zheng, Junkai Xie, Chongli Yuan, Alexander Chubykin, Christopher J Staiger, Guangjun Zhang, Qing Deng

## Abstract

Potassium channels regulate membrane potential and diverse physiological processes, including cell migration. However, the specific function of the inwardly rectifying potassium channels in immune cell chemotaxis is unknown. Here, we identified that the inwardly rectifying potassium channel Kir7.1 (KCNJ13) maintains the resting membrane potential and is required for directional sensing during neutrophil chemotaxis. Pharmacological or genetic inhibition of Kir7.1 in neutrophils impaired direction sensing toward various chemoattractants without affecting cell polarization in multiple neutrophil models. Using genetically encoded voltage indicators, we observed oscillating depolarization of the membrane potential in protrusions in zebrafish neutrophils, and Kir7.1 is required for polarized depolarization towards the chemokine source. Focal depolarization with optogenetic tools biases pseudopod selection and induces *de novo* protrusions. Global hyperpolarizing neutrophils stalled cell migration. Furthermore, Kir7.1 regulates GPCR signaling activation. This work adds membrane potential to the intricate feedforward mechanism, coupling the adaptive and excitable network required to steer immune cells in complex tissue environments.

## Introduction

Neutrophils play a crucial role in innate immunity as the first line of defense against pathogens and critical regulators of the cancer microenvironment (Burn et al., 2021; Zhou et al., 2023). The mobility of neutrophils is essential for their physiological function and is highly regulated. Chemotaxis is one of the best-studied cell migrations, in which the cell moves toward extracellular signaling molecules (chemoattractants), such as cytokines. Cell migration is a coordinated cycle of protrusion generation and tail retraction, primarily regulated by the actin cytoskeleton. Signaling from the G-protein coupled receptors (GPCRs) translates the shallow gradients of the extracellular cues to a steep polarization of intracellular signaling molecules, such as phosphoinositide 3-kinase (PI3K), PI-3-phosphatase PTEN, and monomeric G proteins (Ras, Rho, Rac, Cdc42) to interpret the guidance cues (Hoeller et al., 2016).

Recently, the cell membrane potential, one of the prospects of bioelectricity, was found to regulate the actin cytoskeleton and adhesion signals (Schwab et al., 2012; Schwab et al., 2008). Reciprocally, the actin cytoskeleton and associated proteins regulate ion channel activities, displaying intricate feedback mechanisms (Mazzochi et al., 2006). Membrane potential (MP) across the plasma membrane is a fundamental property of all animal cells that compose a lipid bilayer impermeable to ions (Kulbacka et al., 2017). The transmembrane voltage creates a strong electric field across the thin lipid bilayer, modulating the activity of membrane proteins and solute exchange, which is essential for a broad spectrum of physiological processes, such as the circadian rhythm of neurons and fibroblasts, biological sensing of neurons, and volume control (Abdul Kadir et al., 2018).

Potassium channels are major contributors to establishing cell MP and among the transporter proteins recognized to regulate cell migration (Schwab et al., 2012; Schwab et al., 2008). Potassium and other ion channels are essential for neutrophil recruitment and function (Immler et al., 2018). Previous *ex vivo* measurements using MP dyes indicate that neutrophils’ resting potential is around -74 mV. The chemokines fMLP and C5a induce various degrees of depolarization (Fletcher and Seligmann, 1986; Messerer et al., 2018; Seeds et al., 1985). The dominant current in unstimulated mouse neutrophils composes a constitutively active, external K+-dependent, strong inwardly rectifying current, although the channel was not pinpointed (Masia et al., 2015). The calcium-activated potassium channel Kca3.1 is required for regulated neutrophil volume decrease and chemotaxis without affecting calcium entry or reactive oxygen species production (Henriquez et al., 2016). The voltage-gated potassium channel Kv1.3 supports sustained Ca^2+^ influx and tight adhesion to the blood vessel wall during acute inflammation (Immler et al., 2022). ATP-sensitive K^+^ channels also regulate neutrophil migration (Da Silva-Santos et al., 2002). In addition to K^+^ channels, anion transporters, including voltage-gated chloride/proton antiporter chloride channel-3 (ClC-3) and the swelling-activated Cl-channel (IClswell), are also required for neutrophil transendothelial migration and chemotaxis (Moreland et al., 2006; Volk et al., 2008). However, the spatial-temporal dynamics of the MP during neutrophil migration have not been characterized.

The inwardly rectifying potassium channel subfamily J member 13 (*Kcnj13*, encoding Kir7.1) regulates smooth muscle cytoskeletal organization during mouse tracheal tubulogenesis (Yin et al., 2018). In humans, mutations in the KCNJ13 gene are associated with retinal degeneration, including autosomal-dominant snowflake vitreoretinal degeneration (Hejtmancik et al., 2008), and recessive mutations cause Leber congenital amaurosis, an early-onset form of blindness (Sergouniotis et al., 2011). In zebrafish, various mutations in *kcnj13* are associated with pigment patterns and fin patterning (Podobnik et al., 2020; Silic et al., 2020), suggesting this channel might have a regulatory role in cell migration. The zebrafish is an excellent model for studying neutrophil biology due to the high conservation of genetic, biochemical, morphological, and metalogical properties compared to their mammalian counterparts (Deng and Huttenlocher, 2012).

Here, we took advantage of the zebrafish neutrophil model and investigated the role of Kir7.1 in neutrophil chemotaxis. We observed that Kir7.1 maintains the resting MP and regulates neutrophil directional sensing, with minimal impact on motility. The MP during neutrophil chemotaxis displayed oscillating polarized depolarizations, and depolarization is sufficient to guide neutrophil migration. The elucidation of the specific function of inward rectifying K^+^ channels advanced our knowledge of how bioelectricity regulates the chemotaxis of neutrophils, providing insights into the intricate regulatory network guiding the efficient infiltration of immune cells in complicated tissue environments.

## Results

### Kir regulates neutrophil chemotaxis in zebrafish

Kir7.1 was known to regulate zebrafish pigmentation and fin patterning (Iwashita et al., 2006; Silic et al., 2020). However, its function in neutrophil migration is unknown. We mined our previously published mRNA seq data on sorted zebrafish neutrophils (Hsu et al., 2019) and identified relatively high transcript levels of two Kir channels, *kcnj13* and *kcnj1b*. We performed RT-PCR from sorted neutrophils and confirmed the expression of *kcnj1b*, *kcnj11*, and *kcnj13* (Fig. S1A). We employed VU590, a potent and moderately selective inhibitor for Kir1.1 (encoded by *kcnj1*) (IC50=290 nM) and Kir7.1 (IC50=8 µM) (Weaver and Denton, 2021), to determine where Kir regulates neutrophil migration in 3-day post-fertilization larvae from *Tg(lyzC:mCherry)^pu43^*. VU590 treatment did not alter neutrophil numbers (Fig. 1A) or their spontaneous migration in fish (Fig. 3B, C) but led to a significantly decreased number of neutrophils recruited to tail transection sites (Fig. 1D-F). Neutrophils migrate from caudal hematopoietic tissue (CHT) to the ventral fin fold when the fish is bathed in leukotriene B4 (LTB4), a lipid chemoattractant (Yoo et al., 2011). VU590 treatment significantly reduced neutrophil chemotaxis toward this defined chemoattractant (Fig. 1G-I).

**Figure 1.**
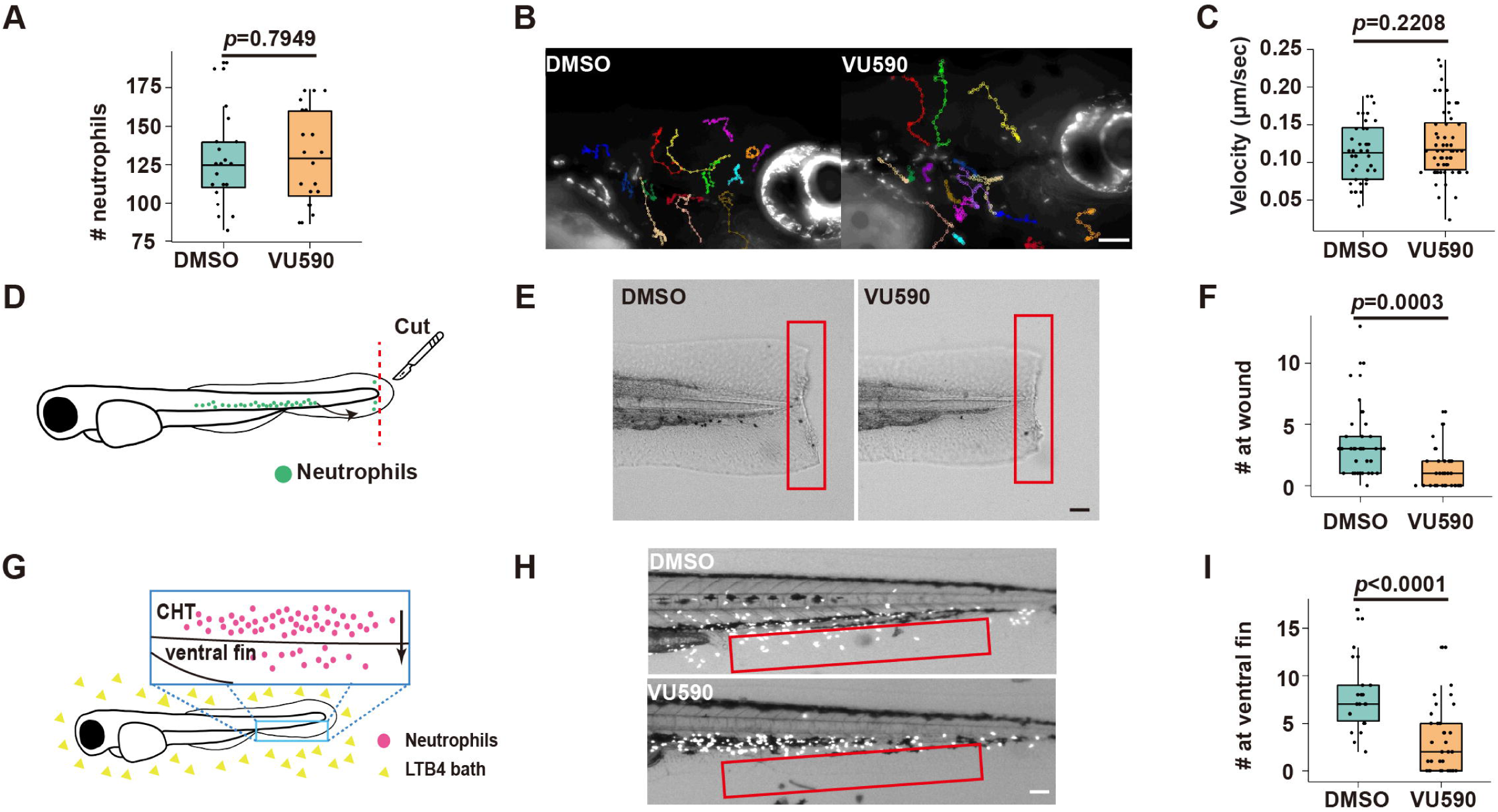
Pharmacological inhibition of Kir7.1 attenuates neutrophil chemotaxis in zebrafish. (A) Numbers of total neutrophils in the zebrafish 3dpf embryos treated with 50 µM VU590 or DMSO. Each dot represents one fish. n > 20 in each group. Data representative of 3 independent experiments. (B) Representative tracks and (C) speed of zebrafish neutrophil spontaneous migration with VU590 or DMSO treatment using *Tg(lyzC:mCherry)pu43*. Scale bar: 50 μm. Each track represents one neutrophil. n >15 neutrophils from 3 fish of each group. (D) Schematic illustration of tail transection and neutrophil recruitment. Green dots represent neutrophils. Neutrophils migrating from caudal hematopoietic tissue (CHT) to the tail 30 min post-injury were counted. (E) Representative images and (F) Quantification of neutrophils recruited to the ventral fin in embryos with VU590 or DMSO treatment. Neutrophils in the boxed regions are quantified. Scale bar: 50 μm. n > 20 in each group. Data representative of 3 independent experiments. (G) Schematic illustration of LTB4-induced neutrophil chemotaxis assay. Red dots represent neutrophils. Neutrophils migrate from caudal hematopoietic tissue (CHT) to the ventral fin in the LTB4 bath. 30 min post-injury were counted. (H) Representative images and quantification (I) for the neutrophils migrated to the ventral fin 30 min in an LTB4 bath with VU590 or DMSO treatment. n > 20 in each group. Data representative of 3 independent experiments. (A, C, F, I) Results are presented as mean ± s.d., Mann–Whitney test.

### Neutrophil intrinsic Kir7.1 regulates chemotaxis in zebrafish

To confirm these findings from the inhibitor, we turned to genetic inhibition. We used a dominant-negative *Jaguar-like* zebrafish (*Kir7.1^G157E^*) that displayed an altered pigment pattern due to a point mutation in the germline (Iwashita et al., 2006; Silic et al., 2020). The neutrophil response to LTB4 or the tail wounds was consistently attenuated in the *Kir7.1^G157E^* fish (Fig. 2A-D). We bred the *Kir7.1^G157E^* line to visualize cell response with a transgenic zebrafish line expressing mCherry in neutrophils. Neutrophils failed to migrate to the ventral fin with a specific directness and directional migration defect but with normal cell velocity (Fig. 2E-H and Movie 1). On the other hand, neutrophil chemotaxis was intact in a *kcnj13* knockout fish line (*kcnj13^pu109^*) (Silic et al., 2020), possibly due to the redundant function of the other Kir family members (Fig. S1B-D).

**Figure 2.**
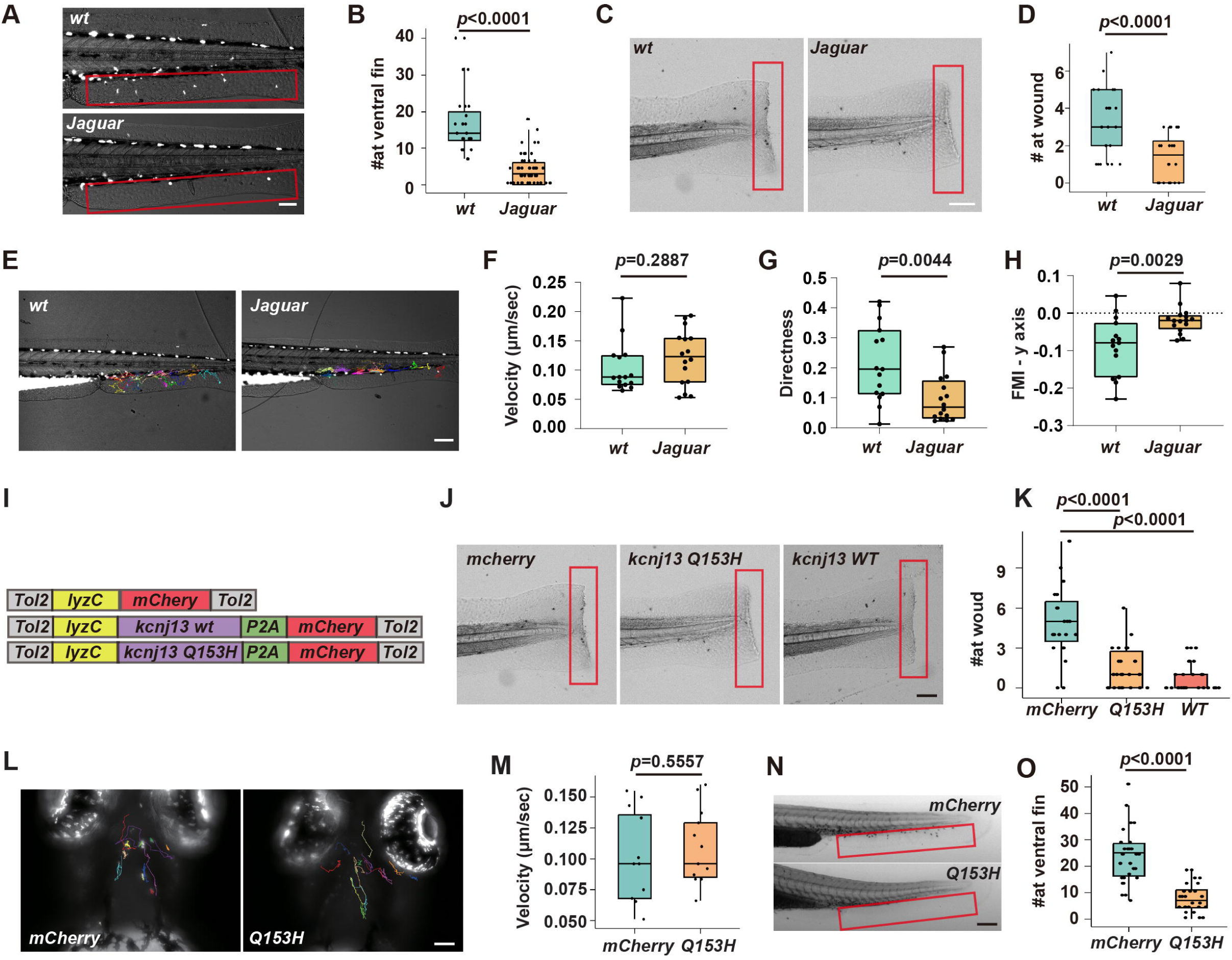
Kir7.1 regulates zebrafish neutrophil chemotaxis. (A) Representative images and quantification (B) for neutrophil response to LTB4 in the *Jaguar* or *wt* sibling controls. Scale bar: 50 μm. n > 20 in each group. Data representative of 3 independent experiments. (C) Representative images and quantification (D) for neutrophil response to tail wounding in the *Jaguar* or *wt* sibling controls. Scale bar: 50 μm. n > 20 in each group. Data representative of 3 independent experiments. (E) Representative tracks of neutrophil migration to the ventral fin in response to LTB4 in the *Jaguar* or *wt* sibling controls. Each track represents one neutrophil. Scale bar: 50 μm. (F-H) Quantification of the speed (F), Directness (G), and forward migration index (H). n > 20 in each group. Data representative of 3 independent experiments. (I) Construct design for neutrophil-specific kcnj13 WT and Q153H mutation overexpression in zebrafish to generate *Tg(lyzC:kcnj13-2A-mCherry)^pu44^*and *Tg(lyzC:kcnj13-Q153H-2A-mCherry)^pu45^*. (J) Representative images and (K) quantifications of neutrophil recruitment to tail fin wound. Scale bar, 300 μm. n > 20 in each group. Data representative of 3 independent experiments. (L) Representative tracks and (M) speed of neutrophils’ spontaneous migration in the head mesenchyme in transgenic lines with neutrophil-specific Kir7.1 Q153H overexpression, *Tg(lyzC:mCherry)^pu35^*expressing mCherry alone is used as a control. Scale bar: 50 μm. n >15 neutrophils from 3 fish of each group. (N) representative images and (O) quantification of the number of neutrophils recruited to the ventral fin upon LTB4 treatment in the indicated zebrafish transgenic lines overexpressing mCherry or Kir7.1Q153H. Scale bar: 50 μm. n > 20 in each group. Data representative of 3 independent experiments. (B, D, F, G, H, M, O) Results are presented as mean ± s.d., Mann–Whitney test. (K) Multiple comparisons, One-way ANOVA.

To disrupt Kir7.1 function specifically in neutrophils, we engineered two transgenic zebrafish lines using a neutrophil-specific lysozyme C (lyzc) promoter (Hsu et al., 2019), *Tg(lyzC:kcnj13-2A-mCherry)^pu44^* and *Tg(lyzC:kcnj13-Q153H-2A-mCherry)^pu45^*, overexpressing wild-type or a dominant-negative version of Kir7.1, Kir7.1^Q153H^ that reduced K^+^ conductance in HEK293T cells when over-expressed (Silic et al., 2020). We co-expressed Kir7.1 with mCherry via a cleavable 2A peptide to visualize neutrophils (Fig. 2I). Neutrophils from *Tg(lyzC:mCherry)^pu43^* expressing mCherry only are used as controls. Wild-type or dominant-negative Kir7.1 overexpression both reduced neutrophil recruitment to wounds, indicating that the channel activity is required in an optimal range to support chemotaxis (Fig. 2J, K). Consistent with the inhibitor treatment, Kir7.1^Q153H^ overexpression significantly reduced neutrophil chemotaxis toward LTB4 without affecting neutrophil random migration (Fig. 2L-O).

### Kir7.1 regulates spatial gradient of plasma membrane potential during neutrophil chemotaxis

To reveal how MP modulates neutrophil chemotaxis, we employed the genetically encoded voltage indicator ASAP3 (Accelerated Sensor of Action Potentials), which was used to monitor action potential in neurons(Villette et al., 2019). We generated a transgenic line *Tg(lyzC: ASAP3)^pu40^* to express ASAP3 specifically in zebrafish neutrophils. We bred this line with *Tg(lyzC: mCherry-CAAX)^pu41^* to label the cell membrane uniformly to mitigate potential artifacts raised by any changes in the membrane thickness. When chemotaxing toward LTB4, neutrophils exhibited a spatial gradient of MP along the axis of movement (Fig. 3A, B, and Movie 2). Cell speed is cyclic, with correlated high speed at both protrusion and tail retraction (Fig. 3C-E). A higher ratio (relatively hyperpolarized) correlates with the retraction speed (Fig. 3F). The protrusion is relatively depolarized, and a slight hyperpolarization also correlates with the reduction in protrusion speed (Fig. 3G). This gradient in MP is cyclic, with the gradient most prominent during active protrusion and absent during tail retraction (Fig.3N). In contrast, this gradient is absent during neutrophil random migration in the head, indicating a response specifically activated by chemokines (Fig. S2A and Movie 3).

**Figure 3.**
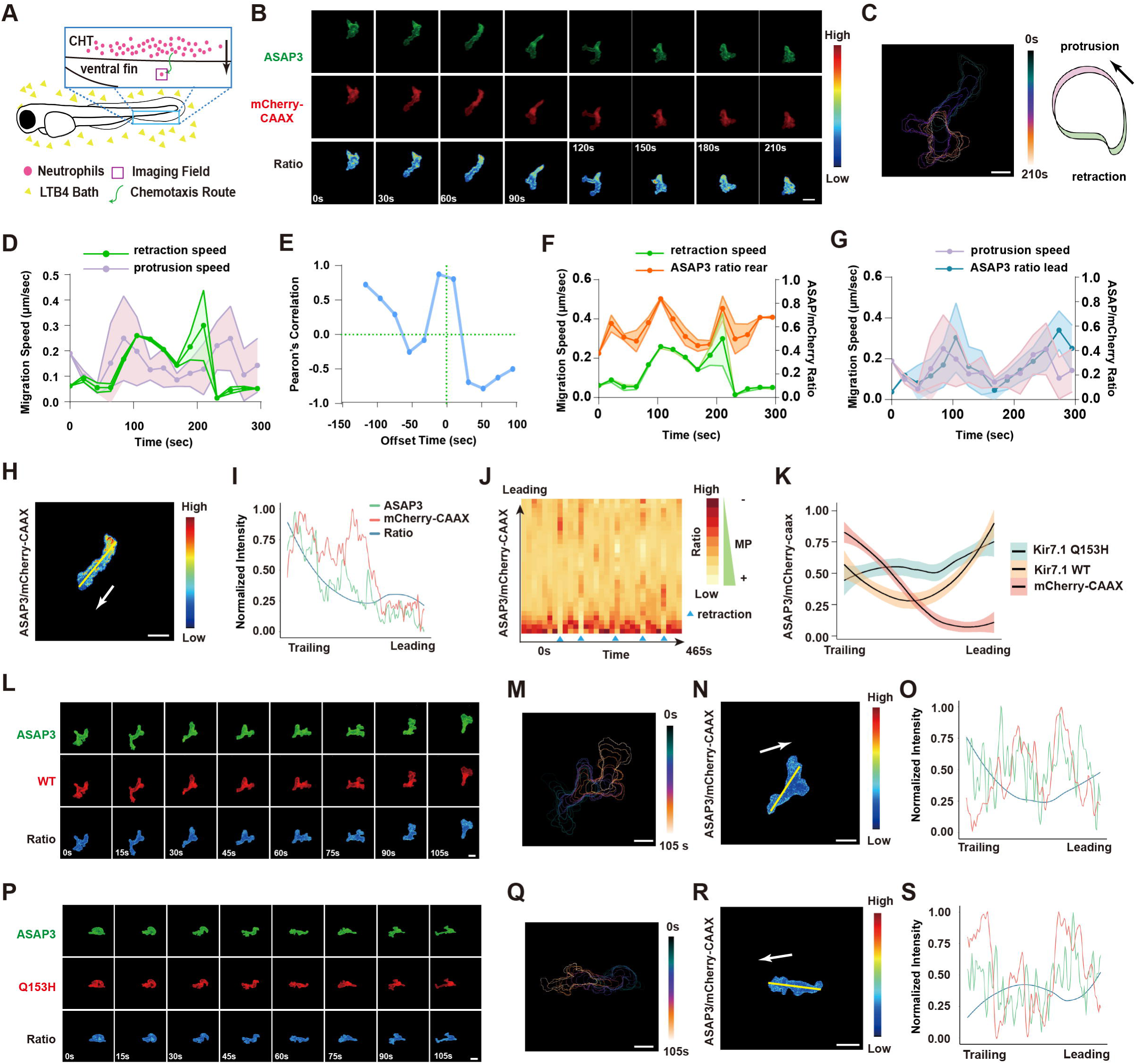
Kir7.1 regulates plasma membrane potential gradient during neutrophil chemotaxis. (A) Schematics of the neutrophils being imaged in the zebrafish ventral fin when migrating toward an LTB4 gradient. (B) Representative time-lapse imaging of ASAP3 and mCherry-CAAX in fish neutrophils migrating towards 30 nM LTB4 using *Tg(lyzC: ASAP3;lyzC: mCherry-CAAX).* Ratiometric analysis (ASAP3/mCherry-CAAX) was performed. (C) Cell outlines over time, indicating cell shape changes during migration. Time was color-coded. Scale bar, 10 μm. Cell outlines were used to quantify cell protrusion (green) and retraction (magenta) speeds. (D) Protrusion (green) and retraction (magenta) speeds for the cell shown in (B). Data is presented as mean ± s.d. for n=3 spots in each group. (E) Pearson’s correlation between the protrusion and retraction speeds for the cell in (B) with indicated time offset. (F, G) ASAP3 ratio and the correlated migration speed at the retraction and the protrusion of cell shown in (B). Data is presented as mean ± s.d. for n=3 spots in each group. (H) A representative frame of the cell in (B) and the ratio of ASAP3/mCherry-CAAX is color-coded. Scale bar, 10μm. (I) Quantification of fluorescence intensity along the indicated line in (H). green: ASAP3; red: mCherry-CAAX; blue: ASAP3/mCherry-CAAX ratio. (J) Kymograph plotting the dynamics of membrane potential gradient change. The time interval between frames is 15 seconds. Blue triangles indicate the tail retraction phase during neutrophil migration. The ratio indicates ASAP3/mCherry-CAAX. MP: membrane potential. (K) Quantitative analysis of the normalized ratio of ASAP3/mCherry along the axis of neutrophils migration upon Kir7.1 WT or Q153H overexpression compared with mCherry-CAAX control. Data is presented as mean ± s.d. for n=15 cells in each group. (L-S) similar analysis of (B, C, H, I) using *Tg(lyzC: ASAP3;lyzC:kcnj13-2A-mCherry-CAAX) or Tg(lyzC: ASAP3;lyzC:kcnj13-Q153H-2A-mCherry-CAAX)*.

To determine the impact of Kir7.1 on neutrophil plasma membrane potential, we generate two additional zebrafish lines, *Tg(lyzC:kcnj13-2A-mCherry-CAAX)^pu46^*and *Tg(lyzC:kcnj13-Q153H-2A-mCherry-CAAX)^pu47^,* overexpressing Kir7.1 wt or Q153H fused with mCherry-CAAX via the P2A peptide. The evident MP depolarization at the leading edge in control neutrophils was lost upon Kir7.1 wt or Q153H overexpression (Fig. 3G). Ratiometric imaging of ASAP3/mCherry-CAAX in neutrophils revealed a significantly attenuated MP gradient upon overexpression of either construct (Fig. 3L-S and movie 4). On the other hand, overall cell polarity was not affected, evidenced by proper localization of the products of the PI3K, Phosphatidylinositol (4,5)-bisphosphate (PIP2), and phosphatidylinositol (3,4,5)-trisphosphate (PIP3), at the leading edge during cell migration (Yoo et al., 2010), albeit not towards the chemokine source (Fig. S2B). Similarly, the intracellular calcium gradient was not altered (Fig. S2C). Our results revealed that Kir7.1 regulates MP gradient during neutrophil chemotaxis, which is likely critical for gradient sensing without affecting neutrophil development, random migration, or cell polarization.

### Plasma membrane depolarization induces protrusions and guides neutrophil migration

We then utilized optogenetic actuators validated in mammalian cells and zebrafish to address how plasma MP regulates neutrophil migration (Antinucci et al., 2020). CoCHR generates high amplitude photocurrent to allow entry of cations, primarily Na^+^ that depolarize cells (Antinucci et al., 2020). We generated a transgenic line *Tg(l*yzC:CoChR-mCherry)^pu42^. To confirm that CoCHR can sufficiently depolarize neutrophils, we illuminated the neutrophils co-expressing ASAP3 and CoChR. ASAP3 signal diminished throughout the cells after focal photo stimulation, which is likely due to the fast spread of the ions outside the initial stimulation region (CoCHR deactivation time is 20 ms, much shorter than the time required for the system to switch from photomanipulation to acquisition mode(Antinucci et al., 2020)) (Fig. 4A, B and Movie 5). Cells quickly recover the MP and continued migration (Fig. 4C). We then determined if local cell membrane depolarization could influence the directional migration or initiate new neutrophil protrusions. When migrating in tissue, neutrophils generate multiple protrusions, and one becomes dominant. We randomly selected one protrusion to stimulate, and 8 out of 10 became dominant and guided cell migration (Fig. 7C and Movie 6). During active cell migration, we targeted one side of the cell body, approximately one-third from the cell front. Small transient protrusions were induced, lasting 30 seconds before retracting back into the cell body in 7 out of 10 cells (Fig.7C and movie 7). Stimulation on the tail did not slow down neutrophil movement (Fig. 7C and Movie 8). This is consistent with previous optogenetics work, which demonstrated that cell polarity is irreversible during persistent migration (Yoo et al., 2010). Our data suggest that membrane depolarization is sufficient to induce protrusion and guide neutrophil migration.

**Figure 4.**
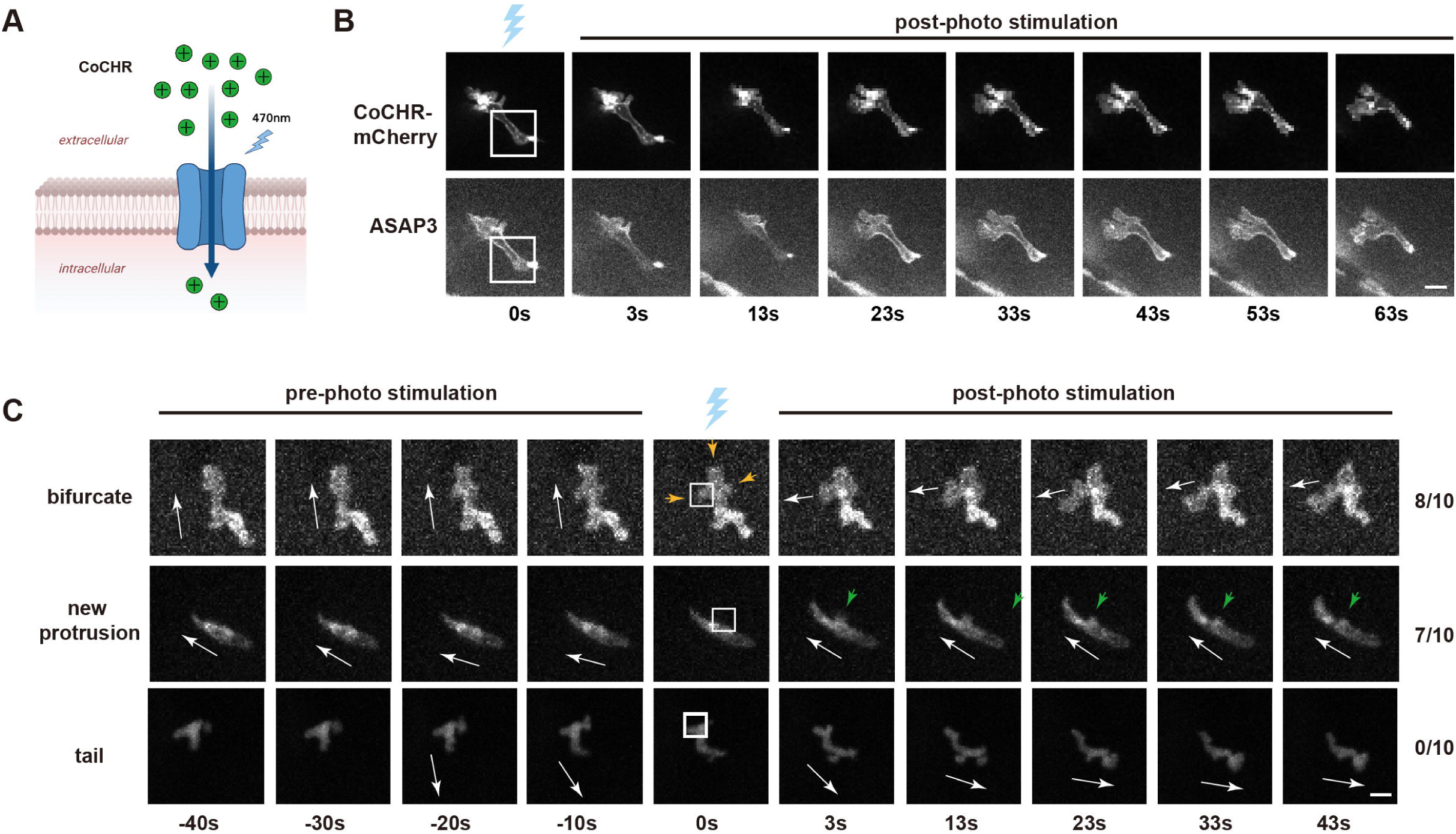
Plasma membrane depolarization induces protrusions and guides neutrophil migration. (A) Schematic of ion selectivity of CoChR. (B) Time lapse of ASAP3 and mCherry after photo stimulation using *Tg(lyzC: ASAP3;lyzC:CoChR-mCherry)^pu42^*. (C) Representative time-lapse images of neutrophil movement before and after the laser stimulation on indicated subcellular areas using *Tg(lyzC:CoChR-mCherry)^pu42^*. The blue laser cartoon indicates the photo-stimulation frame. White arrowheads indicate the direction of cell movement. White boxes indicate the area stimulated. Orange arrowheads represent multiple protrusions. Green arrowheads represent the *de novo* protrusion. Scale bar, 5 μm. N = 10 cells for each condition.

To expand the discovery to other ions, we employed additional optogenetics tools. GtACR2, an anion selective channel, and BLINK2, a K^+^ selective channel, cause hyperpolarization and neuronal activity suppression (Fig. S3A, B). We expressed the actuators in zebrafish neutrophils using transient plasmid injection and exposed the entire cell to blue light (Fig. 2C, D and Movies 9). Cell migration was stalled immediately after light stimulation, which recovered within 2-3 minutes after removing the stimulation. Control cells expressing mCherry were not affected by a similar light stimulation condition. Our results indicate that neutrophil behavior is excitable, reminiscent of the neuronal system.

### Kir regulates GPCR signaling in human neutrophil-like cells

Our data in zebrafish neutrophils indicated a critical role of the MP gradient in neutrophil chemotaxis in an intact tissue environment. Due to the limited number of cells and inaccessibility of the cells in the tissue, we next turned to human neutrophil cell models, differentiated HL-60 cells (dHL-60). Neutrophils adapt their chemotactic behavior based on different extracellular matrix components and respond to various chemoattractants. We performed an under-agarose migration assay on BSA or collagen-coated surfaces and multiple chemoattractants, including fMLP, LTB4, and the complement fragment C5a, to determine if Kir activity is required for general chemotaxis. VU590 reduced neutrophil directionality toward the gradient for all chemoattractants tested, regardless of the substrates (Fig. 5A, B). Notably, the speed of the cells was not significantly different from the DMSO control (Fig. 5C). As a control, we measured cell random migration under agarose. VU590 treatment significantly increased cell velocity; however, in the presence of fMLP (chemokinesis), the velocity was comparable to the DMSO treated control (Fig. 5D). We attempted to express KCNJ13-DN in dHL-60 cells, but did not observe robust phenotype, which is likely due to high expression of other KCNJ family members, such as KCNJ2 and KCNJ15 (Rincon et al., 2018).

**Figure 5.**
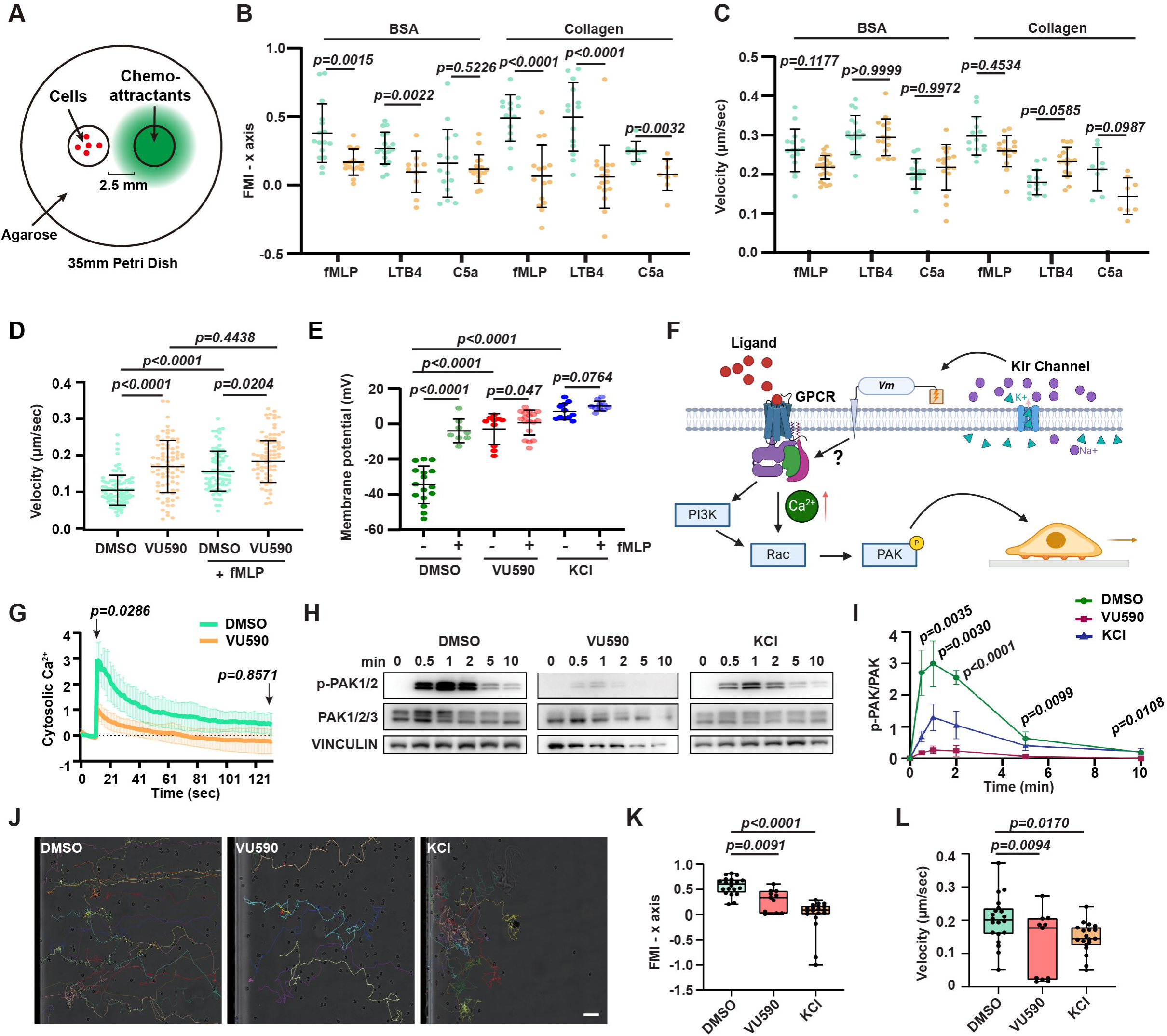
Kir7.1 regulates chemotaxis and GPCR signaling of human neutrophils. (A) Under-agarose migration assay of dHL-60 cells treated with 50 μM VU590 or DMSO on BSA- or Collagen-coated glass bottom dishes towards indicated chemoattractants. (B, C) Quantification of Forward Migration Index (FMI) towards the chemokine gradient and velocity of cells. n > 10 in each group. Data presents mean ± s.d., representative of 3 independent experiments. (D) Under-agarose random migration velocity of differentiated HL-60 cells treated with 50 μM VU590 or DMSO on BSA-coated glass bottom dishes in the presence or absence of uniform fMLP. n =20 in each group. Data presents mean ± s.d., pooled from 3 independent experiments. (E) Whole-cell patch-clamp measurement of differentiated HL-60 cells treated with VU590 or KCl with or without fMLP activation. n > 10 in each group. Data presents mean ± s.d., representative of 3 independent experiments. (F) Illustration of how Kir regulates membrane potential and GPCR signaling. (G) Quantification of intracellular calcium post-fMLP treatment of differentiated HL-60 cells treated with VU590 or DMSO. Data presents mean ± s.d., representative of 3 independent experiments. (H, I) Immunoblot and normalized quantification of time-course PAK activation post-fMLP treatment of differentiated HL-60 cells treated with DMSO, VU590, or KCl. Data presents mean ± s.d. from 3 independent experiments. (J) Representative tracks of PMN chemotaxis toward fMLP after treatment with DMSO, VU590, or KCl, in IBIDI μ-Slide coated with Collagen IV. The lines indicate individual cell trajectories for 120 min. Scale bar: 60 μm. (K, L) Quantification of Forward Migration Index (FMI) towards fMLP and velocity of cells in (L). n > 10 in each group. Data presents mean ± s.d., representative of 3 independent experiments. (B, C, G) Mann–Whitney test. (D, E, I, K, L) Multiple comparisons, One-way ANOVA.

We then measured the MP using a whole cell patch clamp. We also applied 40 mM KCl as a positive control, which alters potassium channel activity and causes cell depolarization (Kilbourne et al., 1991; Rienecker et al., 2020). Differentiated HL-60 cell resting MP was around -30 mV, and fMLP induced a depolarization. VU590 and the positive control KCl depolarized dHL-60 to 0 mV, which does not further depolarize after fMLP stimulation (Fig. 5E). These data suggest that the Kir family maintains the polarized MP in neutrophil-like cells at the resting state. Chemokine binding to chemokine receptors, GPCRs, leads to rapid activation of the downstream signaling pathways, including calcium mobilization from the intracellular store and the p21-activated kinase (PAK) (Itakura et al., 2013), which is specifically localized to the leading edge of migrating neutrophils and drives efficient directional migration (Fig. 5F). The cytoplasmic calcium spike induced by fMLP was significantly blunted upon VU590 treatment (Fig. 5G). Additionally, PAK activation upon fMLP stimulation was attenuated PAK activation in UV590-treated dHL-60 cells (Fig. 5H, I). KCl also induced significant PAK activation suppression, indicating that depolarization is required for optimal GPCR signaling activation.

### Kir regulates membrane potential and chemotaxis of primary human neutrophils

We then validated our results in primary human neutrophils (PMNs). Human peripheral blood neutrophils express 14 members of the Kir family genes, with high levels of *KCNJ2* and *KCNJ15* and intermediate levels of *KCNJ1* and *KCNJ13* (Rincon et al., 2018). In an IBIDI µ-slide chemotaxis assay, consistent with the observation in zebrafish, the VU590 treatment dampened neutrophil chemotaxis towards fMLP, significantly reducing directionality towards the chemokine gradient (Fig. 5J-L and Movie 10). Indeed, KCl treatment also selectively reduced the directionality of neutrophil chemotaxis. Together, the Kir family is necessary for robust gradient sensing in neutrophil chemotaxis in humans and zebrafish.

## Discussion

In this study, we characterized the dynamics of MP in response to chemoattractant stimulation of neutrophils. Our findings reveal that MP and Kir7.1 are crucial in neutrophil chemotaxis. Pharmacological inhibition or genetic alteration of an inward-rectifying potassium ion channel protein abolishes MP gradients and neutrophil chemotaxis without affecting the basal motility in zebrafish. We identified a temporal and spatial gradient in MP along the direction of cell migration during neutrophil chemotaxis, consistent with the rapid shut-off and slower signaling activity adaptation during chemotaxis (Tang et al., 2014). Using cutting-edge optogenetic techniques, we elucidated that locally depolarizing the MP induces *de novo* protrusions, demonstrating a direct cause-and-effect relationship between the neutrophil MP and chemotaxis. On the other hand, prolonged depolarization blunts GPCR signaling activation. These insights open new avenues for exploring the role of different ion channel proteins in leukocyte migration and present potential targets for clinical therapies controlling neutrophil migration and the overall inflammation process.

### Neutrophil membrane potential

The dynamic heterogeneity of MP in neutrophils, visualized by fast-response voltage indicators, is unexpected. All previous work focused on the collective MP in the cell without spatiotemporal resolution. Given the rapid velocity at which electrical signals spread through the cellular structure within submilliseconds, any transient heterogeneity in the distribution of action potential within a small cell was considered too brief to detect. Indeed, the poroelastic structure of the actin meshwork and cytoplasm within the lamellipodium resists water flow, possibly preventing or retarding the immediate equalization of local ion concentrations (Charras et al., 2005; Ito et al., 1992). In addition, it has been proposed that the mitochondrial membrane potentials can be profoundly heterogeneous. Individual crista within the same mitochondrion can have different MP (Smiley et al., 1991) and, thus, potentially function as independent bioenergetic units (Wolf et al., 2019). Even though the MP in mitochondria is produced mainly from proton dynamics, these studies provided that the phospholipid bilayers can preserve the uneven distribution of MP within specific areas for limited periods. In our observations, it is also the case that the MP gradient is prominent in active migration, while dissipated when cells are not actively migrating.

### How Kcnj13 regulates neutrophil chemotaxis

Kir channels’ inwardly rectifying activity is stimulated with hyperpolarization (Xu et al., 2009), consistent with the earlier observation that neutrophil membrane has a high intrinsic permeability to K^+^, which is low or absent in the depolarized state (Majander and Wikstrom, 1989). Our results show that genetic or pharmacological inhibition of Kir7.1 depolarizes neutrophils. A similar observation is made in mouse smooth muscle, whereas Kir7.1 deficiency leads to depolarization and the accumulation of intracellular K^+^ (Yin et al., 2018). Classical Kir and K_ATP_ channel inhibition also depolarizes ventricular myocytes (Hibino et al., 2010). Kir7.1 is a weak inward rectifier carrying measurable outward K^+^ currents at voltages that are positive to the K^+^ reversal potential (Krapivinsky et al., 1998; Xu et al., 2009). Due to the unique channel property, Kir7.1 possibly regulates neutrophil directional sensing via setting the resting MP to maintain excitability, reinforcing local depolarization at the protrusion, or inducing repolarization to allow the cycles of pseudopod selection, similar to the role of Kir in cardiac myocytes (Hibino et al., 2010). It is unlikely that Kir7.1 regulates cell volume, which is associated with overall cell speed (Nagy et al., 2024). It is also worth noting that Kir channel function redundancy in neutrophils as multiple Kir channels are expressed and dominant negative mutants rather than loss of function mutants showed evident migration phenotypes in our study. How the GPCR regulates these Kir channels remains unknown (Ghamari-Langroudi et al., 2015). There is no current evidence that Kir7.1 activity is regulated by GPCR signaling. Our results extend the function of Kir7.1 in immune cells and chemotaxis and provide insights into the intricate Kir7.1-regulated signaling events.

### Membrane potential regulates the excitable network in chemotaxis

It is worth noting that MP regulation is separable from membrane charge (Banerjee et al., 2022) and galvanotaxis (migration in an electrical field), primarily working through the asymmetric distribution of charged molecules. An excitable Ras/PI3K/Rac/F-actin signal transduction network regulates essential cell motility (Sasaki et al., 2007; Tang et al., 2014). Our results prove that restricted local depolarization alone is sufficient for directional guidance. GPCR signaling, primary Gβγ, generates local excitation via Ras/PI3K/PTEN that couples with the actin oscillators to bias cell polarity and directional migration (Hoeller et al., 2016). Indeed, GPCR activities are directly modulated by MP (David et al., 2022; Seifert and Wenzel-Seifert, 2001). Charged residues, hydrogen bonds, and Na^+^ bound to the allosteric site in the transmembrane domains are sensitive to MP, which leads to conformational changes that affect ligand binding affinity or G protein coupling (David et al., 2022). Although the knowledge came from GPCRs in the neuronal or cardiac system, it is likely applicable to chemokine receptors. In addition, voltage-sensing phosphatase is tuned by the gradient of MP and can contribute to phosphoinositide regulation and cell migration (Okamura et al., 2018). MP regulates PIP_2_ localization and Ras signaling (Zhou et al., 2015). Intriguingly, local activation of Rac is sufficient to steer neutrophil migration in tissue, suggesting that the excitable network can be biased without GPCR inputs (Yoo et al., 2010). Our results, therefore, add MP to the intricate feedforward mechanism, coupling the adaptive and excitable networks that mediate eukaryotic chemotaxis. Extensive future work will be needed to pinpoint the exact mechanism.

### GPCR signal leads to local depolarization

Due to the specific local depolarization during cell protrusion, depolarization may be regulated by the GPCR signal, which depolarizes the MP upon stimulation with various ligands (Becker et al., 2024; Lazzari et al., 1990; Messerer et al., 2018; Seligmann et al., 1981). The exact mechanism that leads to the depolarization in protrusion is thus likely a multitude and still needs further investigation. Numerous ion channels directly interact with G proteins or are indirectly modulated by G-protein coupled signaling pathways (Inanobe and Kurachi, 2014), in particular, the Gβγ activates G protein-gated inward rectifying potassium channels (GIRK) (Whorton and MacKinnon, 2013) and a background K^+^ channel TASK-2 (Anazco et al., 2013). In addition, the NADPH complex generates a large amount of H^+^ at the cytosolic face (Fletcher and Seligmann, 1986), possibly contributing to the depolarization upon GPCR activation (Seligmann et al., 1981; Stratmann et al., 2021), although the two can be separated (Seeds et al., 1985; Sullivan et al., 1984). Mechanically activated ion channels (Ranade et al., 2015) are likely involved in setting MP in cell protrusion where the membrane experiences tension.

In *Dictyostelium discoideum*, chemotaxis is intact when plasma MP is depolarized (Van Duijn et al., 1990), indicating a species-specific role of MP in eukaryotic chemotaxis. Meanwhile, MP controls the beating of cirri and reorientates a single-celled eukaryote hypotrich ciliate Euplotes when they swim in fluids or walk on the surface (Laeverenz-Schlogelhofer and Wan, 2024), indicating remarkable evolutionary conservation. In summary, our results provided new evidence of ion channels regulating neutrophil cell migration through Kir mediating directional sensing. Demonstrating the role of bioelectricity in regulating chemokine receptor signaling and directional sensing advances our understanding of the chemotaxis process, which is central to immunology, cancer metastasis, and developmental biology.

## Methods

### Generation of Transgenic and Mutant Zebrafish Lines

The zebrafish experiment was conducted by internationally accepted standards. The Animal Care and Use Protocol was approved by the Purdue Animal Care and Use Committee (PACUC), adhering to the guidelines for using zebrafish in the NIH Intramural Research Program (protocol number: 1401001018). Microinjections of fish embryos were performed by injecting 1nl of a mixture containing 25 ng/µl plasmids and 35 ng/µl tol2 transposase mRNA in an isotonic solution into the cytoplasm of embryos at the one-cell stage. The stable lines were generated as previously described (Deng et al., 2011). At least two founders (F0) for each line were obtained. Experiments were performed using F2 larvae produced by F1 fish derived from multiple founders to minimize the artifacts associated with random insertion sites.

*The Kcnj13* knockout line was generated using gRNAs to target the 6^th^ exon of the *kcnj13* gene, as described previously(Silic et al., 2020). The mutation was confirmed by Sanger sequencing. Before being used for experiments, the mutant fish was outcrossed to the F3 generation.

### Molecular Cloning

A tol2-lyzC vector was used in this study for gene expression in fish neutrophils. Target genes were inserted into the Tol2 backbone containing the lyzC promoter and SV40 polyA for neutrophil-specific expression. Fish *kcnj13* WT and Q153H mutation gene fragments were cloned from the constructs described in our previous studies (Silic et al., 2020). ASAP3 gene was cloned from pAAV-hSyn-ASAP3-WPRE plasmid (addgene #132331); GtACR2 gene was cloned from pTol1-UAS:GtACR2-tdTomato plasmid (addgene #124236); BLINK2 gene was cloned from pDONR-BLINK2 plasmid (addgene #117075); CoCHR gene was cloned from pAAV-Syn-CoChR-GFP plasmid (addgene #59070); PHAKT fragment was cloned from pHR SFFVp PHAkt Cerulean plasmid (addgene #50837) and fused followed by EGFP protein sequence. The mCherry and mCherry-CAAX fragments used in fish and human cells were cloned from our previous plasmid constructs in the lab and inserted into fish and human vectors, respectively.

### Pharmacological Treatment

VU590 (Cayman Chemical, Item No. 15177) was dissolved in dimethyl sulfoxide (DMSO) to make a 100 mM stock and then further diluted in E3 or mHBSS ((modified HBSS with 20mM HEPES and 0.5% FBS)) to a working concentration of 50 μM. DMSO was utilized as vehicle control. For neutrophil recruitment and random motility assays, larvae were pretreated with the inhibitor for 1 h before experimental procedures. KCl was prepared at 2M stock concentration and diluted to a working concentration 40 mM for 1hr pre-treatment on cells. VU590 and KCl working concentrations were maintained throughout the experiment.

### Primary Human Neutrophil Isolation

Primary human neutrophils were obtained from the peripheral blood of healthy adult donors and collected under approval by Purdue University’s Institutional Review Board (IRB). Primary human neutrophils were isolated with Milteny MACSxpress Neutrophil Isolation Kit (Macs # 130-104-434) and RBC lysed for 10 min RT (Thermo # 00-4300-54) as described (Hsu et al., 2022) and subjected to chemotaxis analysis.

### IBIDI µ-slide Chemotaxis

Primary human neutrophils were resuspended in mHBSS (modified HBSS with 20mM HEPES and 0.5% FBS) at 4×10^6^ ml^−1^ and loaded into collagen-coated µ-slides following the manufacturer’s instructions (IBIDI, Cat.No:80322). fMLP was added to the right-hand reservoir at a concentration of 1 µM. Chemotaxis was recorded every 1 min for 2 h using a laser scanning confocal microscope (LSM 710) with a 1.0/10× objective. The tracking and velocity of neutrophils were measured using ImageJ with the MTrackJ plugin and exported for further analysis.

### Under-Agarose Chemotaxis Assay

The under-agarose chemotaxis assay applied in this study was modified from that in a previous study (Heit and Kubes, 2003). Briefly, 0.5% ultrapure agarose gel made from 1% gel in DPBS 1:1 ratio to warm mHBSS was poured into a 35 mm collagen-coated tissue culture petri dish. 3 mL gel was required for each dish. The gel was left at 4°C for 15 min to solidify. Wells were punched by a 1. 5mm diameter metal tube with 0.3mm wall thickness using a customized 3D printing mold. The distance between the two wells designed in the mold is 2 mm. 10^6^-10^7^ pretreated HL-60 cells in mHBSS were loaded into one well and incubated at 37°C for 15 minutes. 500 nM fMLP, 600 nM LTB4, or 10μg/mL C5a in mHBSS were loaded into the adjacent well until the liquid level reached the top. The petri dish was incubated at 37°C for 15 mins, then set up in BioTek Lionheart FX Automated Microscope (Winooski, VT, USA) using a 10× phase objective, Plan Fluorite WD 10 NA 0.3 (1320516) for imaging. Cells were tracked using MTrackJ image J plugin.

### Immunoblotting

dHL-60 cells were differentiated for 6 days in 1.3% DMSO, and 10^6^ cells were pelleted in microcentrifuge tubes. Cells were lysed with RIPA buffer (Thermo #89900), supplemented with a proteinase inhibitor cocktail (Roche # 4693132001). Protein contents in the supernatant were determined with BCA assay (Thermo #23225), and 25∼50 µg of proteins was subjected to SDS-PAGE and transferred to PVDF membranes. Phosho-protein probing was performed as described (Graziano et al., 2017). Briefly, 1.5×106 cells/ml dHL-60s were starved in serum-free media for 1h at 37°C/5% CO2. Cells were then stimulated with 10 nM fMLP for 1 or 3min at RT and immediately put on ice with the addition of ice-cold lysis solution in a 1:1 ratio (20% TCA, 40 mM NaF, and 20 mM β-glycerophosphate (Sigma #T6399, #G9422, #s6776)). Cells were incubated on ice for 1h, and proteins were pelleted down and washed once with 0.75 ml of ice-cold 0.5% TCA and resuspended in 2× Laemmli sample buffer (Bio-Rad Laboratories). Western blot was performed with anti-phospho-PAK1/2 (Cell signaling #2605S), anti-PAK (Cell signaling #2604), and Vinculin as a loading control. Membranes were incubated with primary antibodies O/N at 4°C, then with goat anti-mouse (Invitrogen 35518) or anti-rabbit (Thermo SA5-35571) secondary antibodies. The fluorescence intensity was measured using an Odyssey imaging system (LI-COR). Band intensity was quantified using Image Studio (LI-COR) and normalized with respective loading controls.

### Patch Clamp of Differentiated HL-60 Cells

The neutrophils were centrifuged at 1000g for 5 min in 24-well plates with a surface-treated polystyrene-coated glass slide in each well to make the neutrophils loosely adherent to the glass slide. Filamented borosilicate glass capillaries (BF150-86-10, Sutter Instruments) were pulled by a micropipette puller (P-1000, Sutter Instruments) for whole-cell patch-clamp recording electrodes to a resistance of 4-6 MΩ. The glass electrodes were filled with an internal solution containing 20 mM KCl, 100 mM K-gluconate, 10 mM HEPES, 4 mM MgATP, 0.3 mM Na2GTP, and 7 mM phosphocreatine. The pH was adjusted to 7.4, and the osmolarity was adjusted to 300 mOsm. The cultured neutrophils on the glass slide were transferred to the recording chamber perfused with a modified Tyrode bath solution (150 mM NaCl, 4 mM KCl, 2 mM MgCl2, 10 mM glucose, 10 mM HEPES, and 2 mM CaCl2 [pH 7.4, 310 mOsm]) (Chubykin et al., 2007) at 32 °C. fMLP, VU590, and 10 M KCl were added to the Tyrode bath accordingly. The whole-cell patching was conducted using open-source software, Autopatcher IG, developed previously (Wu and Chubykin, 2017; Wu, et al., 2016). The patch clamp recording signals were processed with an amplifier (Multiclamp 700B, Molecular Devices) and a digitizer (Digitata, 1550, Molecular Devices). Acquired data were collected using Clampex software (Axon Instruments, Union City, CA) with a 10,000-Hz low-pass filter. Continuous potential recordings by the current clamp were filtered at 2 kHz. The resting MP was calculated by averaging 0-1 sec continuous potential measurement.

### RT-PCR

Total RNA of neutrophils sorted from adult kidney marrow (Hsu et al., 2019) was extracted using a RNeasy Plus Mini Kit (no. 74104; Qiagen). RT-PCR was performed with QIAGEN OneStep RT-PCR Kit (210212). The following primers were used: *dre-ef1a+:* 5′-TACGCCTGGGTGTTGGACAAA-3′;

*dre-ef1a-:*5′-TCTTCTTGATGTATCCGCTGA-3′; *dre-mpx+*: 5′-ACCAGTGAGCCTGAGACACGCA-3′; *dre-mpx-*: 5′- TGCAGACACCGCTGGCAGTT-3′; *dre-krt4+*: 5′- CTATGGAAGTGGTCTTGGTGGAGG-3′; *dre-krt4-*: 5′- CCTGAAGAGCATCAACCTTGGC-3′;

*dre-kcnj1a1+*: 5′- CTCCACACTGAAGAAAGAGCTTGCT-3′; *dre-kcnj1a1-*: 5′- ATTTTGCCACAACAGGTCGCC-3′; *dre-kcnj1b+*: 5′-GGTTACGTGTTTCTCCCCGT-3 ′; *dre-kcnj1b-*: 5′-ACAAGCAGCTCGCTAGGTTT-3′; *dre-kcnj2b+*: 5′- ATCGGCTTGAGCTTTGACCAAC-3′; *dre-kcnj2b-*: 5′-CTTTTCCATTGCCGTAGCCGT-3 ′; *dre-kcnj10a+*: 5′-TCGCAGACTAAAGAGGGGGA-3′; *dre-kcnj10a-*: 5′- TCCATCAGAGCCCCAGGTTA-3′; *dre-kcnj11+*: 5′-TTCGTCGCCAAAAACGGAAC-3′ ; *dre-kcnj11-*: 5′-GTGGACGGTGGCACTGATAA-3′; *dre-kcnj13+*: 5′- AGGCCTTCATCACTGGTGC-3′; *dre-kcnj13-*: 5′-TGGATCTCGTCAGGCAGGTA-3′.

### Tailfin wounding and Sudan Black staining

Tailfin wounding and Sudan Black staining were carried out with embryos at 3 days post fertilization (dpf) as described previously (Deng et al., 2011). Briefly, embryos were fixed in 4% paraformaldehyde in phosphate-buffered saline overnight at 4°C and stained with Sudan Black.

### Fluorescent Confocal Microscopy

Fluorescent microscopy imaging data were obtained with a Nikon A1R fluorescent confocal microscope. Larvae at 3 dpf were settled on a glass-bottom dish, and imaging was performed at 28 °C. Neutrophil random motility imaging was described before (Deng et al., 2011).30 nM LTB4 was introduced to fish in E3 to induce neutrophil chemotaxis(Yoo et al., 2011). Time-lapse images for zebrafish neutrophil random migration and LTB4-induced chemotaxis were acquired using a 20× objective lens at 1 min intervals for 30 mins. Biosensor imaging was acquired by a 60× oil objective lens. The green and red channels were acquired simultaneously with a 488-nm and 561-nm laser, respectively. Zebrafish neutrophils with biosensors were scanned and imaged by the z-stack function at 15s intervals for 5 min.

### Optogenetics Manipulation

Spinning-disk confocal microscopy (SDCM) was performed using a Yokogawa scanner unit CSU-X1-A1 mounted on an Olympus IX-83 microscope equipped with a 40X 1.0–numerical aperture (NA) UPlanApo oil immersion objective (Olympus) and an Andor iXon Ultra 897BV EMCCD camera (Andor Technology). mCherry fluorescence was excited with the 561-nm laser line, and emission was collected through a 610/37-nm filter. Time-lapse images were collected at 10-s intervals to track the individual neutrophils, and z-series at 2-µm step size for eight steps were acquired at each time point. The photoactivation of optogenetic reporters was performed with an Andor Mosaic3 photostimulation module (Oxford Instruments) integrated into the SDCM system. For photoactivation, the time series was paused, an ROI was drawn on the interested part of the neutrophil, and a 445-nm (1.3 W) laser with 50% power and 3s exposure was used to irradiate the ROI.

### Cytoplasmic Calcium Measurements

Cytosolic Ca^2+^ was measured using Fluo-4 Ca^2+^ Imaging Kit (F10489, Invitrogen). 1ml of dHL-60 cells at 5 x 10^5^ cells/ml were treated with 50 µM VU590 or DMSO control at 37°C for 2.5h. The cells were then incubated with PowerLoad solution and Fluo-4 dye at 37°C for 15 min and then at room temperature for another 15 min. After incubation, cells were washed with mHBSS twice and resuspended in 1ml mHBSS at 5 x 10^5^ cells/ml. The resuspension was then treated again with DMSO or VU590 inhibitor. 200 µL of cells were loaded into fibrinogen-coated µ-Slide 8 Well High (80806, IBIDI) and left to settle at 37°C for 30 min before imaging. Green fluorescence images were recorded with a BioTek Lionheart FX Automated Microscope (Winooski, VT, USA) with 20 × phase lens at 1 s interval for 10 s to obtain baseline fluorescence. 20 µl of 1 µM fMLP was then added into wells using a dispenser, and images were recorded for another 2 min with 1 s interval. The resulting mean fluorescence intensity was normalized to that of the initial 10 s before fMLP addition.

### Image Data Analysis

Image analysis was performed using Imaris software and Fiji/ImageJ. For zebrafish fish neutrophil fluorescent image data, all images at the same time point of a single cell were projected to 2D for analysis. The cell images were segmented, the background was subtracted in Imaris, and the processed images were imported to Fiji/ImageJ. Temporal color-coded cell outlines were plotted as described (Pal et al., 2023). Three spots were manually selected at both the leading and trailing edges of each neutrophil across all time points. The tracks for each spot over time were calculated using the built-in Spots function in Imaris software. Protrusion and retraction speeds were then exported through the same function. The rolling Pearson’s correlation was calculated with a five-time-frame window as previously described (Tsai et al., 2019). Line scans were performed in Fiji/ImageJ by drawing a straight-line segment inside the cells with a line of 12 pixels to obtain an average intensity value. The values of biosensor fluorescent intensity were divided by that of mCherry or mCherry-CAAX to perform ratiometric analysis. The ratio or individual channel intensity profiles were smoothened using the Savitzky– Golay method and plotted in R. The kymograph was also plotted in R/Rstudio after gathering all time points’ biosensor/loading control normalized (min-max method) ratioed intensity.

### Statistical Analysis

Statistical analysis was carried out using PRISM 6 (GraphPad). Mann–Whitney test and two-tailed Student’s t-test were used to determine the statistical significance of differences between two groups, and One-way ANOVA was used to determine the statistical significance of differences between the groups above two. A P-value of less than 0.05 was considered statistically significant.

## Author Contribution

GZ and DQ conceived the project. TW, DK, CD, DHW, WA, MS, WZ, KS, XC, JX, and TK performed experiments. TW, CD, and TK prepared the figures. AC, CS, GZ, CY, and DQ supervised the project. TW, WZ, XC, DQ, and GZ wrote the manuscript.

## Acknowledgment

We thank Dr. Xiaoguang Zhu for his guidance on microscopes at the Bindley Bioscience Center, a core facility of the NIH-funded Indiana Clinical and Translational Sciences Institute. The HL-60 clonal cell line is a gift from Dr. Orion Weiner (University of California, San Francisco). The *Jaguar-* like fish was a gift from Dr. David M. Parichy (University of Virginia). The work was supported by research funding from the National Institutes of Health (R35GM119787 to QD, R35GM-124913 to GZ) and (P30CA023168 to Purdue Center for Cancer Research) for shared resources. This work is based upon efforts supported by EMBRIO Institute, contract #2120200, a National Science Foundation (NSF) Biology Integration Institute.

## Competing interests

The authors declare no competing financial or non-financial interests.

## Notes

### Competing Interest Statement

The authors have declared no competing interest.

