## Supplementary material for "Inwardly rectifying potassium channels regulate membrane potential polarization and direction sensing during neutrophil chemotaxis": Movie legend

**Movie 1. Point mutation in *kcnj13* decreases zebrafish neutrophil chemotaxis by LTB4.**

Time-lapse imaging of zebrafish neutrophil chemotaxis towards LTB4 in the ventral fin of the *Jaguar* mutant and the *wt* sibling control. The image started immediately after the LTB4 addition. The migration trajectory is displayed over the tracked cell. Scale bar: 200 μm.

**Movie 2. The polarity of membrane potential in zebrafish neutrophil chemotaxis.**

Ratiometric imaging of ASAP3 and mCherry-CAAX during zebrafish neutrophil chemotaxis towards LTB4 in the ventral fin in *Tg(lyzC: ASAP3;lyzC: mCherry-CAAX)*. The heatmap indicates a high-to-low ratio of ASAP3/mCherry. Scale bar: 10 μm.

**Movie 3. The polarity of membrane potential in zebrafish neutrophil random migration.**

Ratiometric imaging of ASAP3 and mCherry-CAAX during neutrophil spontaneous migration in the head mesenchyme in *Tg(lyzC: ASAP3;lyzC: mCherry-CAAX)*. The heatmap indicates a high-to-low ratio of ASAP3/mCherry. Scale bar: 10 μm.

**Movie 4. The polarity of membrane potential in zebrafish neutrophil chemotaxis upon Kir7.1 overexpression.**

Ratiometric imaging of ASAP3 and mCherry-CAAX during zebrafish neutrophil chemotaxis toward LTB4 in the ventral fin in *Tg(lyzC: ASAP3;lyzC:kcnj13-2A-mCherry-CAAX)*(left) or *Tg(lyzC: ASAP3;lyzC:kcnj13-Q153H-2A-mCherry-CAAX)* (right). The heatmap indicates a high-to-low ratio of ASAP3/mCherry. Scale bar: 10 μm.

**Movie 5. Photo-stimulation effectively regulates zebrafish neutrophil membrane potential.**

Time-lapse imaging of zebrafish neutrophil spontaneous migration in the caudal hematopoietic tissue before and after light stimulation. Neutrophils overexpress ASAP3 and CoCHR-mCherry-CAAX in *Tg(lyzC: ASAP3;lyzC:CoChR-mCherry*. Box indicates the frame and region of stimulation. Scale bar: 10 μm.

**Movie 6. Photo-stimulation biases pseudopod selection and neutrophil migration.**

Time-lapse imaging of zebrafish neutrophil spontaneous migration in the head mesenchyme before and after light stimulation in *Tg(lyzC: CoChR-mCherry)^pu42^*. Box indicates the frame and the pseudopod for stimulation. Scale bar: 10 μm.

**Movie 7. Photo-stimulation induces de novo protrusion during neutrophil migration.**

Time-lapse imaging of zebrafish neutrophil spontaneous migration in the head mesenchyme before and after light stimulation in *Tg(lyzC: CoChR-mCherry)^pu42^*. Box indicates the frame and the pseudopod for stimulation. Scale bar: 10 μm.

**Movie 8. Photo-stimulation at the tail of zebrafish neutrophils cannot reverse directionality.**

Time-lapse imaging of zebrafish neutrophil spontaneous migration in the head mesenchyme before and after light stimulation in *Tg(lyzC: CoChR-mCherry)^pu42^*. Box indicates the frame and the pseudopod for stimulation. Scale bar: 10 μm.

**Movies 9. Photo-stimulation during neutrophil spontaneous migration when cells overexpress GtARC2, BLINK2, or the mCherry control.**

Time-lapse imaging of zebrafish neutrophil spontaneous migration in the head mesenchyme before and after light stimulation. Neutrophils overexpress GtARC2-mCherry, BLINK2-P2A-mCherry, or the mCherry control. Box indicates the frame and region of stimulation. White asterisks indicate the time frames of the migration halt. Scale bar: 10 μm.

**Movie 10. Pharmacological inhibition of Kir7.1 during primary human neutrophil chemotaxis towards fMLP.**

Time-lapse imaging of human neutrophil chemotaxis towards fMLP in an IBIDI chemotaxis chamber treated with DMSO, VU590, or KCl. The migration trajectory is displayed over the tracked cell. Scale bar: 10 μm.
