## Supplementary Figure for "Inwardly rectifying potassium channels regulate membrane potential polarization and direction sensing during neutrophil chemotaxis"

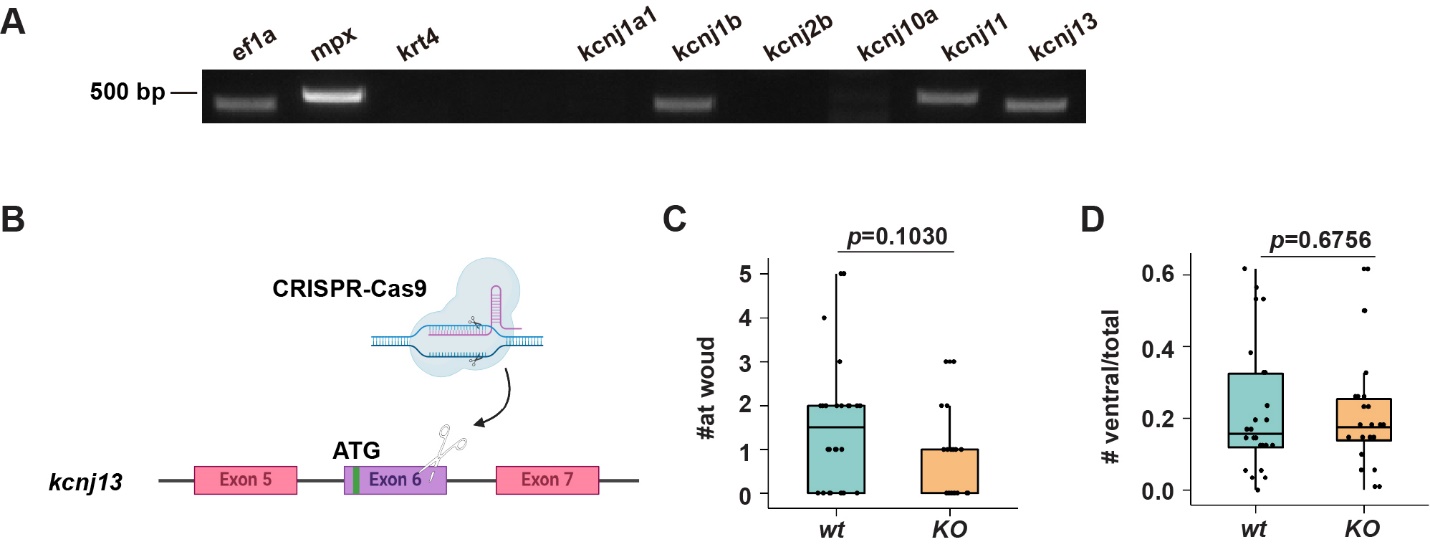

**Figure S1. Kir7.1 regulates zebrafish neutrophil chemotaxis.** (A) RT-PCR of indicated genes using mRNA extracted from FACS-sorted neutrophils. (B) Schematic of the *kncj13* knockout fish. (C, D) quantifications of neutrophil recruited to the tail wound (C) or ventral fin induced by LTB4 (D) in kcnj13 KO fish. (C, D) Each dot represents one neutrophil. n > 20 in each group. Data representative of 3 independent experiments. Results are presented as mean ± s.d., Mann–Whitney test.

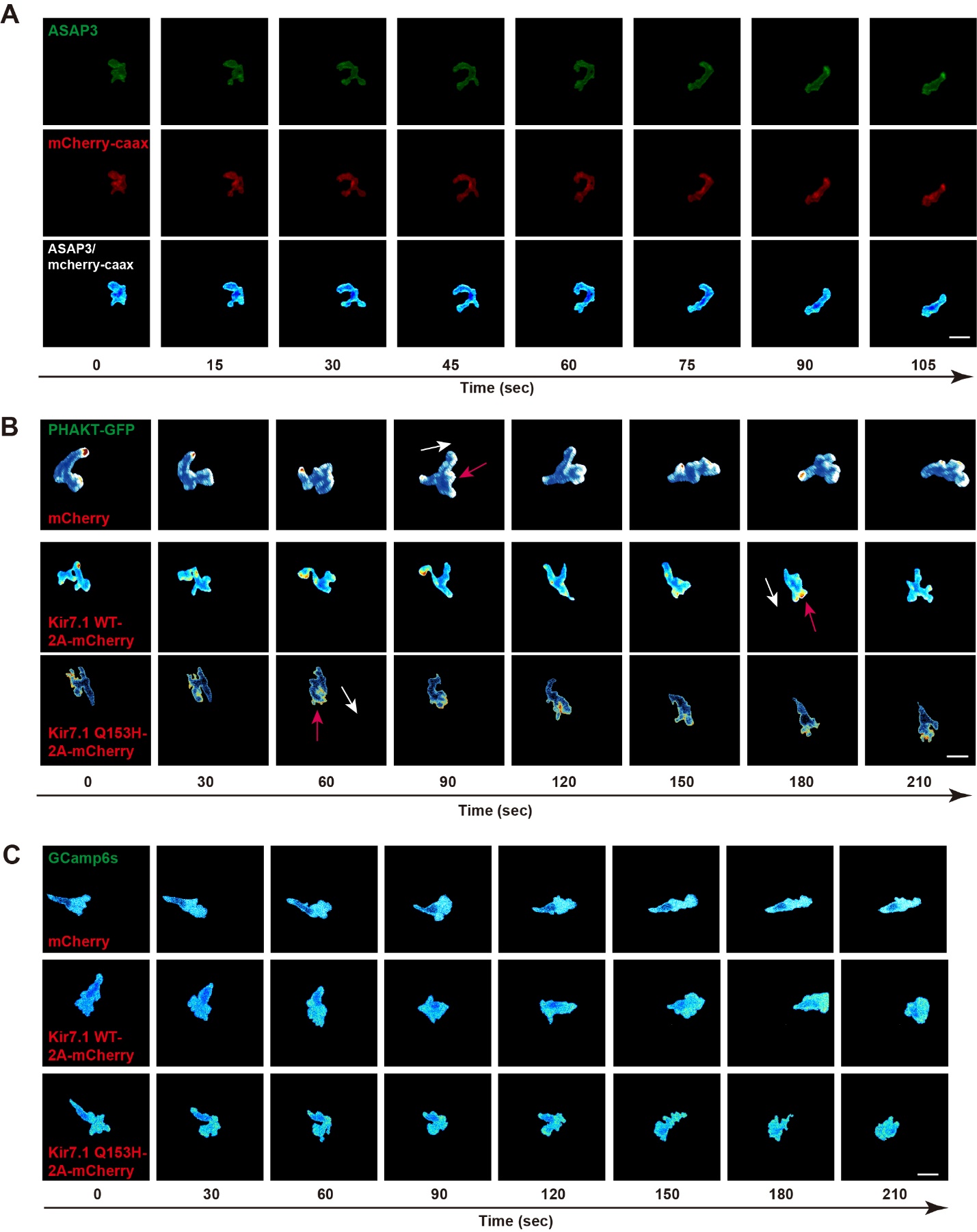

**Figure S2. Kir7.1 does not regulate cell polarity.** (A) Representative time-lapse imaging of ASAP3 and mCherry-CAAX in fish neutrophils randomly migrating in the head mesenchyme using *Tg(lyzC: ASAP3;lyzC: mCherry-CAAX).* Ratiometric analysis (ASAP3/mCherry-CAAX) was performed. Scale bars: 10 μm. Image representatives of 15 neutrophils. (B) Representative ratiometric images of PHAKT-GFP and mCherry during neutrophils migration to LTB4 in zebrafish *Tg(lyzC:mCherry;lyzC:PHART-GFP)*, *Tg(lyzC:kcnj13-2A-mCherry;lyzC:PHART-GFP)* or *Tg(lyzC:kcnj13-Q153H-2A-mCherry;lyzC:PHAKT-GFP)*. The ratio indicates PHAKT-GFP/mCherry. White arrows indicate the direction of cell migration. Red arrows point to protrusions with a high accumulation of PHAKT. Scale bars: 10 μm. Image representatives of 15 neutrophils from each group. (C) Representative ratiometric images of GCamp6s and mCherry during neutrophils migration to LTB4 in zebrafish *Tg(lyzC:mCherry;lyzC:GCamp6s)*, *Tg(lyzC:kcnj13-2A-mCherry;lyzC: GCamp6s)* or *Tg(lyzC:kcnj13-Q153H-2A-mCherry;lyzC: GCamp6s)*. The ratio indicates GCamp6S/mCherry. White arrows indicate the direction of cell migration. Red arrows point to areas with high calcium flux. Scale bars: 10 μm. Image representatives of 15 neutrophils from each group.

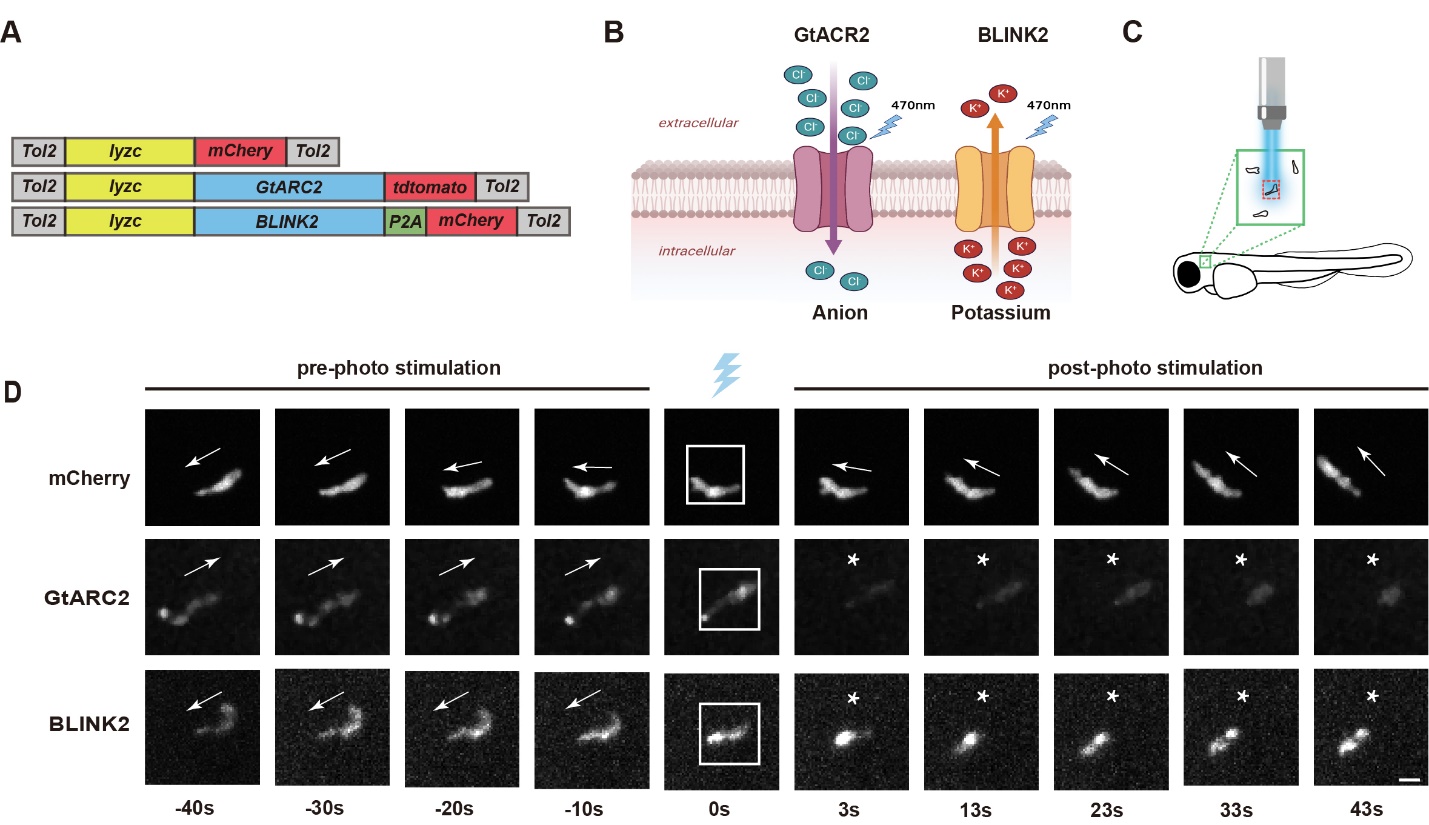

**Figure S3. Global Plasma membrane depolarization stalls neutrophil migration.** (A, B) Schematic of the construct design and ion selectivity of the optogenetic actuators. (C) Illustration of the neutrophils being imaged in the head mesenchyme. (D) Representative time-lapse images of neutrophil movement under the control of different optogenetic actuators or the mCherry control before and after stimulating using a 445 nm laser. The blue laser cartoon indicates the photo-stimulation frame. White Box indicates the stimulation area. White asterisks indicate the time frames of halted migration. Scale bar, 10 μm. N=20 for each optogenetic manipulation group.
